# Identification of B56α, B56γ and BAP1 as PRR14L binding partners

**DOI:** 10.1101/2023.04.29.538809

**Authors:** Andrew Chase, Gonzalo Carreño-Tarragona, Feng Lin, Sarah Yapp, Joanna Score, Catherine Bryant, Nicholas CP Cross

**Author notes:** Correspondence to: Professor N.C.P. Cross, Wessex Genomics Laboratory Service, Salisbury NHS Foundation Trust, Salisbury SP2 8BJ, UK.

## Abstract

Truncating mutations have been previously described in *PRR14L* associated with acquired isodisomy of chromosome 22 in myeloid neoplasms. Very little is known about the function of PRR14L, but previous work showed localization to the midbody and binding to KIF4A. Here we confirm binding of PRR14L to PP2A components B56α and B56γ. Similar to the related protein PRR14, PRR14L binds B56α via a conserved short linear motif within the C-terminal Tantalus domain. We also confirmed binding to BAP1, which forms the H2A deubiquinating complex PR-DUB with ASXL1, thereby linking PRR14L to a protein with established leukemogenic significance. AlphaFold data predicts PRR14L structure to be largely disordered, consistent with a possible role as a scaffold protein.

## Introduction

The development of large-scale genomic sequencing and associated technologies has given us an almost complete picture of the genetic changes that lead to hemopoietic neoplasms. An understanding of the pathogenic mechanisms driven by many of these mutated genes, however, is lacking. *PRR14L* is one such gene; inactivating mutations, usually associated with acquired isodisomy of chromosome 22, have been found in chronic myelomonocytic leukemia (CMML), related myeloid neoplasms and age-related clonal hematopoiesis, but very little is known about PRR14L’s function. We previously found that knockdown of *PRR14L* in human CD34+ cells lead to an increase in monocytes and decrease in granulocytes in differentiation assays, consistent with the finding of mutations in CMML. We also confirmed KIF4A as a binding partner with likely localization to the midbody.

Here we show that PRR14L binds to members of the PP2A complex and BAP1. BAP1 forms a complex with ASXL1, a gene also mutated in myeloid neoplasms including CMML, thereby providing a link to a protein with known leukemogenic activity. Using CRISPR-Cas9 technology we have inserted FLAG tags into endogenous *PRR14L* in HEK293 cells, which may be useful for further proteomic studies. Consistent with the lack of obvious protein domains outside of the C-terminal Tantalus domain, we discuss evidence for PRR14L being an intrinsically disordered protein.

## Methods

### Cell lines

The HEK293F cell line (Thermo Fisher Scientific, Waltham, MA, USA) was grown in DMEM plus 10% fetal calf serum and transfected using Lipofectamine 2000 (Invitrogen, Carlsbad, CA, USA).

### Expression constructs

C-terminal Myc-FLAG tagged *PRR14L* in pCMV6 was previously described (Chase et al., 2019). The following *PRR14L* constructs were also created: (i) PRR14L-R1833X, with introduction of a stop codon at amino acid R1833 to recreate a deletion mutant identified in a patient with CMML and acquired chromosome 22 isodisomy; (ii) mutation of amino acids 2067-8 from EE to AA, a mutation previously shown to abolish binding of B56α at the PRR14 short linear motif (SLiM) (Dunlevy et al., 2020); (iii) PRR14L-TANT, a fragment of *PRR14L* consisting of the C-terminal 143 amino acids (amino acids 2009-2151) containing the Tantalus domain and SLiM sequence by PCR amplification of the required coding region with insertion into pCMV6-AC-Myc-DDK (Origene, Rockville, MD, USA) using an In-Fusion HD Cloning Plus kit (Takara Bio, Shiga, Japan); (iv) PRR14L-Q314X, a fragment of *PRR14L* consisting of the N-terminal 313 amino acids by mutagenesis of codon 314 of Myc-FLAG N-terminal tagged *PRR14L* to a stop codon. This was designed since N-terminal PRR14L shows some conservation within vertebrates and may have retained conserved protein-binding function.

V245 pCEP-4HA B56α (Seeling et al., 1999) was a gift from David Virshup (Addgene plasmid # 14532 ; http://n2t.net/addgene:14532 ; RRID:Addgene_14532).

To create HA-tagged expression constructs for putative PRR14L binding partners, the coding sequences of *PPP2R5C* (B56γ) and *CIT* were amplified from HEK293 cDNA. Both were inserted into pCMV6-AN-HA (Origene, #PS100013) and the correct sequence confirmed by sequencing. The inserts correspond to references sequences NM_001352913.2 (*PPP2R5C*) and XM_017018737.1 (*CIT*, identical to NM_001206999.2 but with serine 1962 removed by alternative splicing).

pcDNA3.1-BAP1 (NM_004656.4) was purchase from Genscript (#OHu20549D) and transferred to pCMV-AN-HA.

### Immunoprecipitation

FLAG tagged WT or mutagenized PRR14L and HA-tagged potential binding protein were transfected together into HEK293 cells. At 48 hours cells protein lysates were prepared (lysis buffer: 10 mM Tris-HCl pH 7.5, 150 mM NaCl, 0.5 mM EDTA and 0.5% NP-40 plus protease inhibitors). Lysates were rolled at 4C for 1 hour with anti-FLAG M2 agarose beads, then washed (wash buffer: 10 mM Tris-HCl ph 7.5, 500 mM NaCl, 0.5 mM EDTA plus protease inhibitors).

The following antibodies were used for western blotting: PRR14L (Origene, Rockville, MD; TA331394), HA (Proteintech, Illinois, USA; 51064-2-AP), Rabbit anti-FLAG (Signalway Antibody LLC, Maryland, USA; T503-2), tubulin (Abcam, Cambridge, UK; ab7291)

### Introduction of a FLAG tag into endogenous *PRR14L* in HEK293F cells by CRISPR-Cas9

3xFLAG was introduced into the N-terminus of *PRR14L* in HEK-293 cells using methodology developed by the Doyon laboratory (Dalvai et al., 2015). hCas9 (Mali et al., 2013) was a gift from George Church (Addgene plasmid # 41815 ; http://n2t.net/addgene:41815 ; RRID:Addgene_41815). MLM3636 was a gift from Keith Joung (Addgene plasmid # 43860 ; http://n2t.net/addgene:43860 ; RRID:Addgene_43860). AAVS1_Puro_PGK1_3xFLAG_Twin_Strep (Dalvai et al., 2015) was a gift from Yannick Doyon (Addgene plasmid # 68375 ; http://n2t.net/addgene:68375; RRID:Addgene_68375).

Cloning of the guide constructs was as described by Keith Joung (supplementary document accompanying Addgene plasmid MLM3636 at https://www.addgene.org/43860/). The guide was created by amplifying approximately 300 bp fragments upstream and downstream of the start codon. Primers for the downstream fragment (left arm) were 5’-gtgacatatgAAATTGAGAATTGTTCATAAGGCAGTATTTG and 5’-ctatccatggtggctaggacagaTGCAGATTATG. Primers to amplify the upstream fragment (right arm) were 5’-ctagttcgaaaagggcgccctgtcatcTGGAGTAGAGAC and 5’-gactgaattcACCTCCTGCTGTGGAGTCCACTAG. These inserted a KOZAK site adjacent to the ATG start codon which was later deleted by mutagenesis to leave the endogenous translation initiation site.

To insert the FLAG tag into endogenous *PRR14L* in HEK293F cells, the cells were transfected with hCas9, MLM3636, and the guide. Seven days after transfection, individual clones were isolated and screened for the FLAG insertion by PCR, a larger band size indicating a possible FLAG insertion. Two positive clones, A3 and D2, with N-terminal insertion of 3xFLAG into endogenous PRR14L, were confirmed by sequencing and western blot.

### Disorder prediction analysis

AlphaFold structural prediction data was accessed at https://alphafold.ebi.ac.uk/. pLDDT data is stored in the B-factor field of the protein PDB file. DisCoP was accessed at http://biomine.cs.vcu.edu/servers/disCoP/, flDPnn at http://biomine.cs.vcu.edu/servers/flDPnn/ and FoldIndex(C) at https://fold.proteopedia.org/cgi-bin/findex.

Protein alignments and similarity calculations were performed using EMBOSS-water, accessed at https://www.ebi.ac.uk/Tools/psa/emboss_water

## Results

### PRR14L binds to BAP1 and PP2A proteins B56α and B56γ

Previous studies using proteomic methodologies have identified potential binding partners of PRR14L. These include several members of the PP2A complex: PPP2R1A (Glatter et al., 2009; Hein et al., 2015; Huttlin et al., 2021; Yadav et al., 2017), PPP2CA (Glatter et al., 2009; Yadav et al., 2017), PPP2CB (Glatter et al., 2009; Huttlin et al., 2015, 2017, 2021) and PPP2R5E (Huttlin et al., 2021). In addition, PPP2R5A (B56α) was recently identified as a binding partner of PRR14, a protein which shows homology with PRR14L within the C-terminal Tantalus domain (Dunlevy et al., 2020). We performed immunoprecipitation assays with B56α and B56γ and demonstrated binding of PRR14L to both proteins (Figure 1A, B).

**Figure 1.**
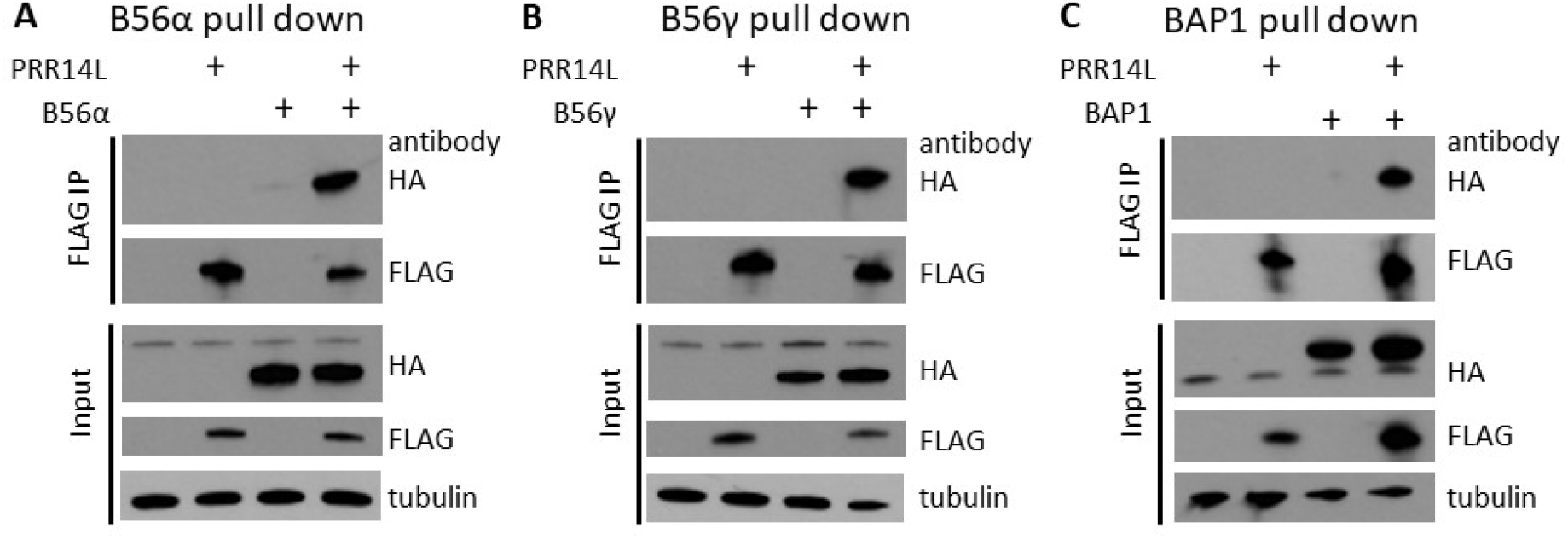
PRR14L binds to B56α, B56γ and BAP1. FLAG-tagged PRR14L and HA-tagged potential binding partners were co-transfected into HEK293 cells and immunoprecipitated with mouse anti-FLAG beads. Pulled down proteins were detected with anti-HA and rabbit anti-FLAG antibodies. We were able to demonstrate binding of B56α (A), B56γ (B) and BAP1 (C) to PRR14L.

BAP1 has also been identified as a potential PRR14L binding protein in a proteomic study (Hein et al, 2015) and is of interest since BAP1 is known to form a complex with ASXL1 which is frequently mutated in myeloid neoplasms (Asada et al., 2018). Using immunoprecipitation we were also able to show binding of BAP1 to PRR14L (Figure 1C), thereby providing a direct link to a protein already known to have leukemogenic activity.

### B56α and B56γ, but not BAP1, bind via the PRR14L SLiM sequence

Dunlevy et al (2020) identified a SLiM sequence (FETIFEE) within the Tantalus domain of PRR14 that mediates binding to PPP2R5A. Mutagenesis of key residues was associated with loss of binding. We mutated the PRR14L SLiM sequence from LETIFEE to LETIFAA (amino acids 2067-2068) and similarly found loss of binding (Figure 2A).

**Figure 2:**
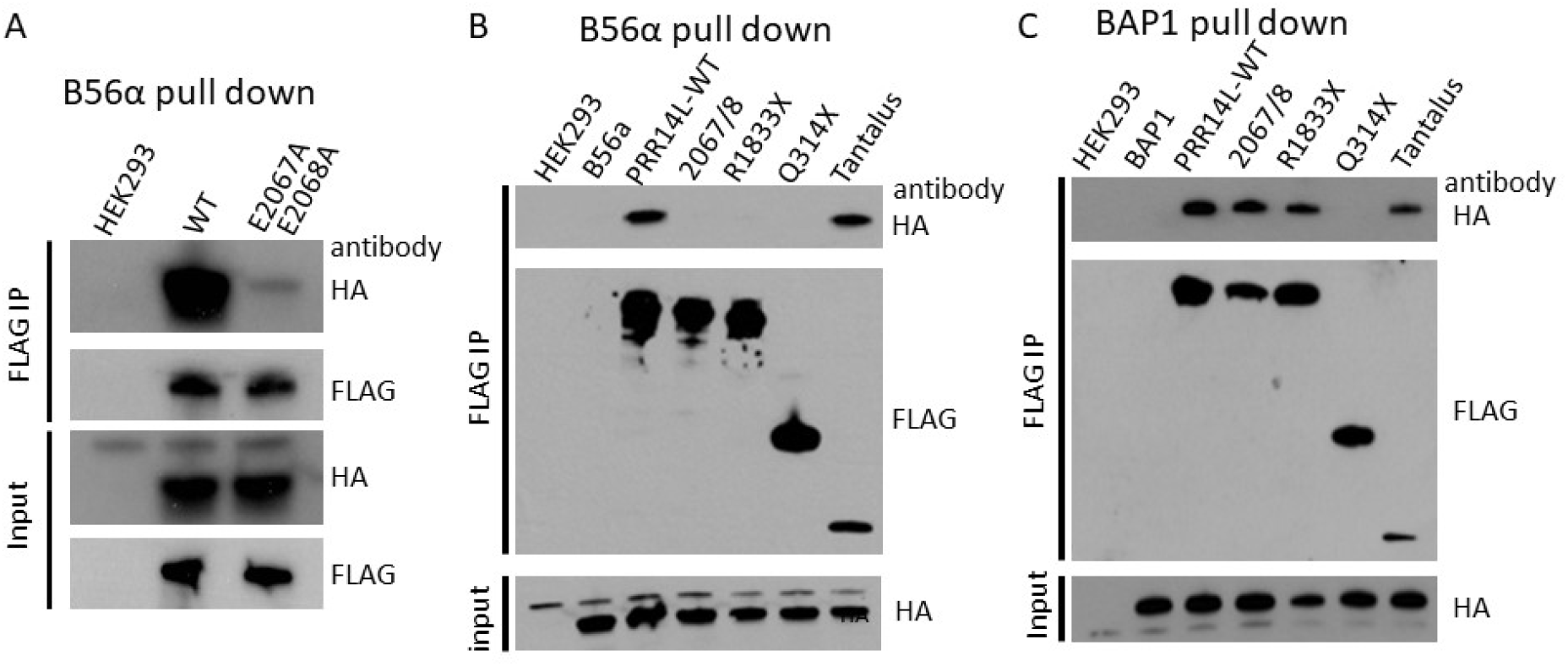
PRR14L with a mutated SLiM sequence and PRR14L fragments were used to identify regions required for binding by B56α and BAP1. Mutation of the PRR14L SLiM amino acids 2067-2068 EE>AA resulted in loss of B56 α binding (A). IP with a series of PRR14L deletion mutants confirmed that only the C-terminal tantalus homology, SLiM-containing region binds B56α (B). In contrast, BAP1 was pulled down by all fragments, including PRR14L with a mutated SLiM sequence, suggesting that BAP1 is bound directly and/or indirectly by two regions AA 314-8133 and the Tantalus homology region AAs 2009-2151 (C).

BAP1 is not known to bind SLiM sequences, and to further delineate PRR14L binding domains in PP2A proteins and BAP1 we created three PRR14L deletion constructs, PRR14L-TANT, PRR14L-Q314X and PRR14L-R1833. As expected, B56α was only pulled down by full length PRR14L and the Tantalus domain (Figure 2B). In contrast, BAP1 was pulled down by all constructs except for PRR14L-Q314X. This included PRR14L-2067/8EE>AA showing that BAP1 binding of PRR14L is not via the SLiM sequence (Figure 2C).

Proteomic data has identified PRR14L as a potential binding partner with CIT (Citron Rho-Interacting Serine/Threonine Kinase) (Capalbo et al., 2019), which localizes to the central spindle and midbody (Bassi et al., 2013) along with KIF4A during cell division. We also performed co-transfection and co-immunoprecipitation experiments with PRR14L-FLAG and CIT-HA, but were unable to demonstrate an interaction.

### PRR14L may be an intrinsically disordered protein

There are currently no experimental structural studies of PRR14L. The artificial intelligence protein structure prediction software AlphaFold (Jumper et al., 2021; Varadi et al., 2022) has been used to predict protein structures for all human proteins including PRR14L. One per-residue measure of confidence provided by AlphaFold is the pLDDT (predicted local distance difference test) score (0-100, 100 being high confidence); values of >70 can be regarded as good or high accuracy whilst values of less than 50 cannot be interpreted and are considered to be strong predictors of disorder. For PRR14L, 92% of residues have pLDDT values <50 (Figure 3) suggesting that most of PRR14L is disordered. Two C-terminal regions show pLDDT values >70; residues 1613-1625 and residues between 2025-2144, the latter region containing the Tantalus domain.

**Figure 3:**
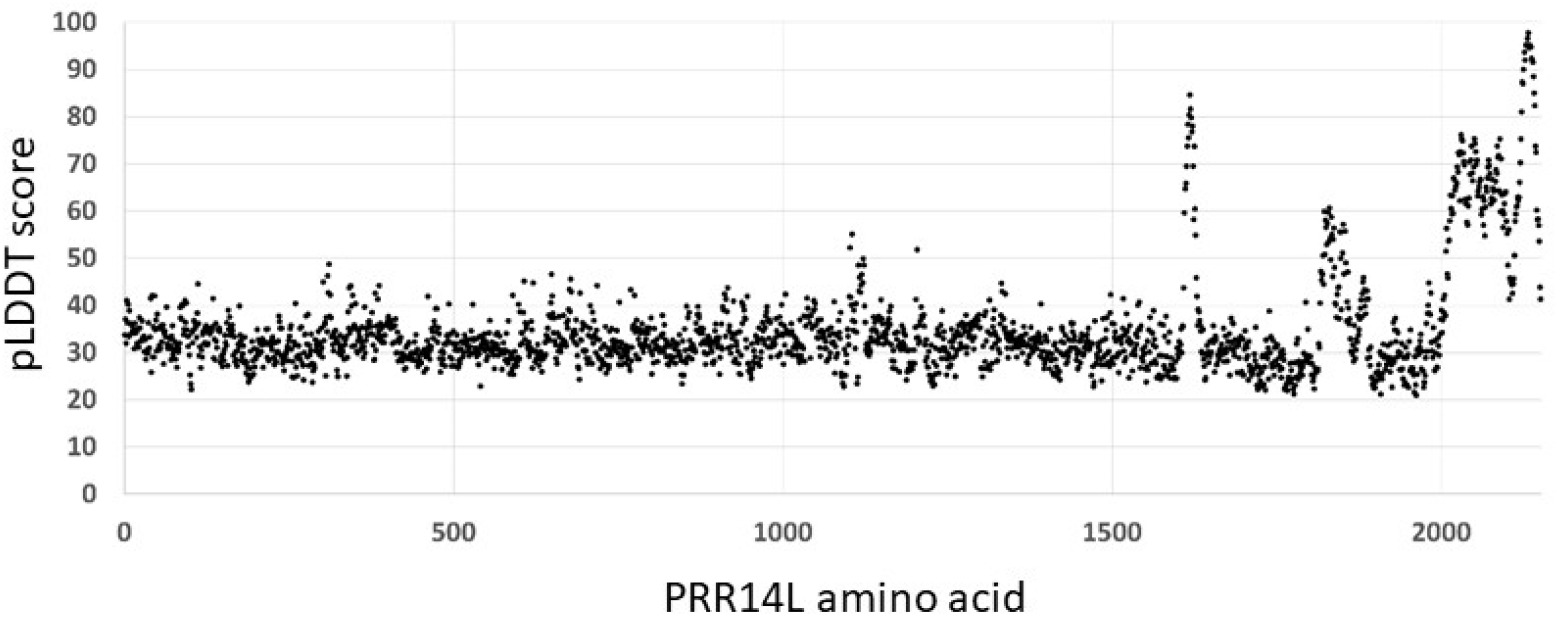
pLDDT scores were obtained from the AlphaFold PDB file. Scores of <50 are considered to be good indicators of disorder, and therefore suggest that PRR14L is largely disordered.

Other *in silico* disorder prediction methods were contradictory, e.g. FoldIndex(C) predicted most of PRR14L to be unfolded and DisCoP predicted a similar pattern of disorder to AlphaFold, whilst flDPnn predicted most of PRR14L to have an ordered structure. Further experimental approaches, e.g. nuclear magnetic resonance (Jensen et al., 2013), may be required to establish the structure and disordered status of PRR14L.

A higher rate of evolutionary change has been described for disordered sequences compared with ordered sequences (Brown et al., 2002), perhaps because disordered regions have fewer structural constraints. We noticed less conservation with other species outside of the C-terminal Tantalus domain region than within the region. We divided PRR14L into N- and C-terminal regions (AAs 1-1609 and 1610-2151) and identified the homologous amino acid in eight vertebrate species.

Similarity between the C- and N-terminal regions in human PRR14L compared with other vertebrates was calculated using EMBOSS-water. We found evolutionary change to be clearly higher outside of the Tantalus region, consistent with the majority of PRR14L being disordered (Table 1).

**Table 1:**
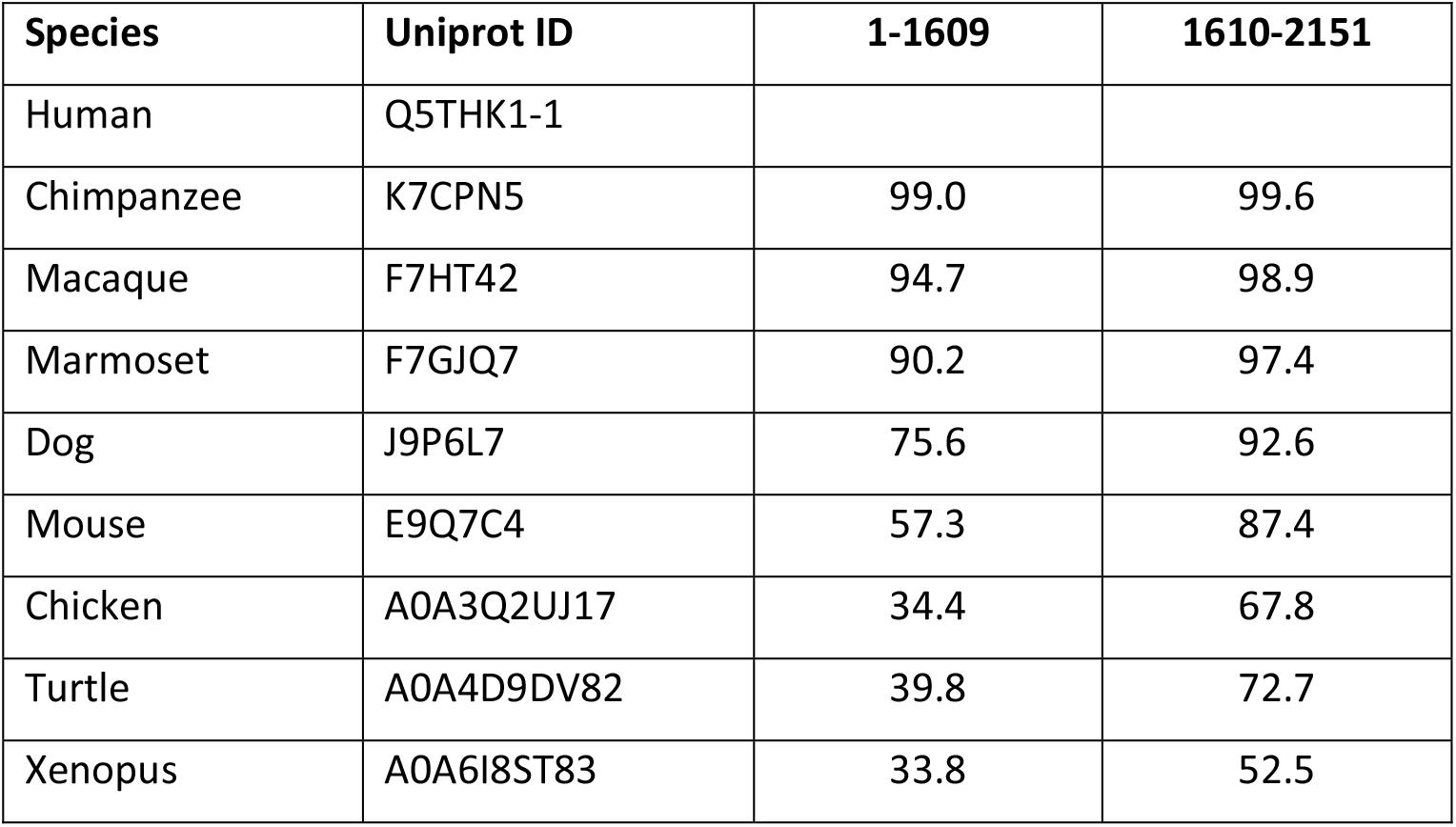
PRR14L protein sequence divergence within vertebrates. The EMBOSS-water tool was used to align protein sequences and calculate similarity. Regions homologous to human PRR14L N- and C-terminal regions 1-1609 and 1610-2151 were identified for several vertebrate species. Sequence divergence was much higher outside of the conserved C-terminal Tantalus homology region (1610-2151), consistent with the bulk of PRR14L being intrinsically disordered.

## Discussion

We previously described inactivating mutations of *PRR14L* associated with chromosome 22 acquired isodisomy, related neoplasms and age-related clonal hematopoiesis (Chase et al., 2019). Using immunofluorescence we showed likely localization of PRR14L to the midbody during cell division.

Two other findings were consistent with midbody localization; we showed an interaction of PRR14L and the midbody protein KIF4A by co-immunoprecipitation, and identified dysregulation of Gαi signaling in CFU-GM after PRR14L knockdown. Gαi proteins are located in the midbody and centrosomes, and reduced expression of Gαi proteins can result in mitotic defects. We also knocked down expression of PRR14L in human CD34+ cells and found an increase in monocytes and a decrease in neutrophils after *in vitro* growth and differentiation assays, consistent with the finding of mutations in CMML and related disorders.

Here we describe further work to investigate other potential PRR14L binding partners. Several members of the PP2A complex had been identified as potential PRR14L-binders in mass-spectrometry based proteomic studies (Glatter et al., 2009; Hein et al., 2015; Huttlin et al., 2015, 2017, 2021; Yadav et al., 2017), and B56α has been shown to bind the related protein PRR14 (Dunlevy et al., 2020). Since B56α and B56γ are reported to be localized primarily in the cytoplasm and nucleus respectively (McCright et al., 1996; Varadkar et al., 2017), and because B56γ, but not B56α, interacts with KIF4A at the anaphase central spindle (Bastos et al., 2014), we investigated interaction with both B56α and B56γ. Both proteins were found to interact with PRR14L by co-immunoprecipitation. Both PRR14 and PRR14L have SLiM sequences within the C-terminal region with homology to Drosophila Tantalus (Dunlevy et al., 2020). To confirm that the PRR14L SLiM mediates binding (as has been shown for PRR14), we mutated the SLiM sequence from LETIFEE to LETIFAA and found that mutation of the SLiM sequence also abolished B56α binding. The significance of PP2A binding by PRR14L is not known, but B56 proteins interact with KIF4 at the anaphase spindle (Bastos et al., 2014) and it is therefore possible that PRR14L interacts with PP2A and KIF4A during the mitotic process. The SliM sequence lies within the Tantalus domain and will be lost by all but one of the patient mutations we previously identified. In the exception, a frameshift lies 27 amino acids C-terminal of the SLiM which may compromise function. Loss or compromised PP2A binding may therefore contribute to the pathogenic activity of mutant PRR14L.

Proteomic studies have also identified interactions between BAP1 and PRR14L (Hein et al., 2015). We were previously unable to pull down endogenous BAP1 with endogenous or tagged PRR14L (Chase et al., 2019), but using co-transfection with HA-tagged BAP1 and FLAG-tagged PRR14L we were able to show interaction by co-immunoprecipitation. BAP1 is not known to bind SLiM sequences and to investigate the interaction between PRR14L and BAP1 further, we performed co-immunoprecipitation experiments with BAP1 and PRR14L with wildtype and mutated SLiM sequences, and PRR14L fragments (PRR14L-R1833X; PRR14L-Q314X; PRR14L-TANT, AAs 2009-2151). In contrast to B56α, interaction by co-immunoprecipitation was found for all PRR14L fragments except for the N-terminal fragment (AAs 1-313). This suggests that binding of BAP1 is not via the SLiM sequence and that two regions, AAs 314-1832, and AAs 2009-2151 containing the Tantalus domain, both directly and/or indirectly interact with BAP1. Since BAP1, but not B56α, is pulled down by PRR14L with a mutated SLiM, BAP1 binding does not require binding of B56α.

The interaction with BAP1, a chromatin deubiquitinating enzyme, is of interest since aberrant activation of BAP1 is thought to be leukemogenic. ASXL1 and BAP1 form part of a H2A deubiquitinase complex PR-DUB, and C-terminal truncation mutations of ASXL1 stabilize the complex and increase BAP1 deubiquitinase activity thereby promoting a leukemogenic gene expression signature (Asada et al., 2018; Sahtoe et al., 2016; Wang et al., 2021). Future work to assess the effect of loss of PRR14L on ASXL1/BAP1 activity would be of interest.

AlphaFold data suggests that the majority of PRR14L is disordered. SLiM sequences are particularly prevalent in disordered proteins and are important mediators of binding between disordered proteins and their partners. The related protein PRR14 has been shown to mediate tethering of heterochromatin to the nuclear lamina (Dunlevy et al., 2020) and the authors suggested that PRR14L, although unlikely to have a tethering function, may act as a scaffold protein. Indeed, a common function of intrinsically disordered proteins is to act a mediators of complex formation (Uversky, 2015) including those involving signaling pathways.

In summary, our findings confirm the interaction of PRR14L with PP2A components and BAP1, thus providing a potential link between signalling pathways and chromatin modifications in clonal disorders associated with *PRR14L* mutations.

